# Instant Automated Inference of Perceived Mental Stress through Smartphone PPG and Thermal Imaging

**DOI:** 10.1101/326157

**Authors:** Youngjun Cho, Simon J. Julier, Nadia Bianchi-Berthouze

## Abstract

**Background:** A smartphone is a promising tool for daily cardiovascular measurement and mental stress monitoring. Photoplethysmography (PPG) and low-cost thermography can be used to create cheap, convenient and mobile systems. However, to achieve robustness, a person has to remain still for several minutes while a measurement is being taken. This is very cumbersome, and limits the usage in applications such producing instant measurements of stress.

**Objective:** We propose to use smartphone-based mobile PPG and thermal imaging to provide a fast binary measure of stress responses to an event using dynamical physiological changes which occur within 20 seconds of the event finishing.

**Methods:** We propose a system that uses a smartphone and its physiological sensors to reliably and continuously measure over a short window of time a person’s blood volume pulse, the time interval between heartbeats (R-R interval) and the 1D thermal signature of the nose tip. 17 healthy participants, involved in a series of stress-inducing mental activities, measured their physiological response to stress in the 20 second-window immediately following each activity. A 10-cm Visual Analogue Scale was used by them to self-report their level of mental stress. As a main labeling strategy, normalized K-means clustering is used to better treat interpersonal differences in ratings. By taking an array of the R-R intervals and thermal directionality as a low-level feature input, we mainly use an artificial neural network to enable the automatic feature learning and the machine learning inference process. To compare the automated inference performance, we also extracted widely used high level features from HRV (e.g., LF/HF ratio) and the thermal signature and input them to a k-nearest neighbor to infer perceived stress levels.

**Results:** First, we tested the physiological measurement reliability. The measured cardiac signals were considered highly reliable (signal goodness probability used, Mean=0.9584, SD=0.0151). The proposed 1D thermal signal processing algorithm effectively minimized the effect of respiratory cycles on detecting the apparent temperature of the nose tip (respiratory signal goodness probability Mean=0.8998 to Mean=0). Second, we tested the 20 seconds instant perceived stress inference performance. The best results were obtained by using automatic feature learning and classification using artificial neural networks rather than using pre-crafted features. The combination of both modalities produced higher accuracy on the binary classification task using 17-fold leave-one-subject-out (LOSO) cross-validation (accuracy: HRV+Thermal: 76.96%; HRV: 60.29%; Thermal: 61.37%). The results are comparable with the state of the art automatic stress recognition methods requiring long term measurements (a minimum of 2 minutes for up to around 80% accuracy from LOSO). Lastly, we explored the impact of different data labeling strategies used in the field on the sensitivity of our inference methods and the need for normalization within individual.

**Conclusions:** Results demonstrate the capability of smartphone biomedical imaging in instant mental stress recognition. Given that this approach does not require long measurements requiring attention and reduced mobility, it is more feasible for mobile mental healthcare solution in the wild.

## Introduction

Human physiological events are controlled by the activities of the sympathetic nervous system (SNS) and parasympathetic nervous system (PNS). Of the many different types, cardiovascular and respiratory events are of importance for the monitoring of a person’s mental health and stress [1–5]. Recent studies on physiological measurements using mobile, low-cost biomedical imaging have shown that it is possible to use a smartphone RGB camera to accurately measure blood volume pulse (BVP) [6–8] and a mobile thermal camera to measure respiratory cycles [9]. Given the advanced mobile biomedical imaging capabilities fully integrated into a smartphone, the smartphone could be a powerful apparatus to monitor and support a person’s mental stress management on a daily basis.

Heart rate variability (HRV), a time series of variation in blood volume pulse (BVP), has been shown to be an informative physiological signature representing the sympathovagal balance between the SNS and PNS [4,10–12]. This can be of use in quantifying a person’s mental stress given that the fight-or-flight responses are a reaction of the autonomic nervous system to stressors, leading to alternations of accelerated and decelerated cardiovascular patterns due to the SNS and PNS, respectively [1,13].

Hand-crafting features from R-R interval sequences (or inter-beat interval, NN-interval) have been explored to quantify HRV and mental stress [4,10,11,14]. Amongst the features, some statistical features (e.g., standard deviation) and frequency-band features (e.g., the normalized power in a frequency band of interest) have been widely used. Empirical findings show that the low frequency band (LF) between 0.04 Hz and 0.15 Hz is likely to be influenced by the SNS and the high frequency band (HF) between 0.15 Hz and 0.4Hz is likely to be affected by the PNS [10]. Therefore, the LF/HR ratio has been used as a stress indicator in many studies [4,11,15,16]. On the other hand, there has been a criticism that this parameter has been used to directly represents stress levels, ignoring the fact that a person’s physiological event cannot be adequately summarized with a single numeric value [17,18]. Therefore, multiple HRV features have been used together with a variety of features from other physiological activities such as perspiratory and respiratory activities for automatically inferring perceived mental stress (e.g., [19,20]) and stressful contexts (e.g., driving tasks [21], desk activities [22]). Empirically, a sliding window of a minimum of 2 minutes window in [17,22] has been considered necessary to extract a number of reliable and informative features to enable machine learning classifiers to perform properly.

Another documented cardiovascular event which happens as a reaction to mental stressors is vasoconstriction of blood vessels over a person’s nasal peripheral tissues [23,24]. This causes a drop in blood flow, results in a decrease in temperature which can be detected by monitoring the temperature of the nose tip [25]. An earlier study using a contact-based multi-channel thermistor revealed that, compared with forehead temperature, there a significant decrease in temperature over the nasal area happens due to mental stress [26]. The same result has been repeatedly reported from the use of thermal imaging in mental stress induction studies [23,27]. However, this approach requires the manual extraction of the temperature from facial thermal images before the stress event occurs, and compares it with the temperature during or after the stress event has finished. This may limit its real-life applications. The automation of the measurement process may improve its use cases. In particular, recent advanced thermal ROI tracking algorithms could help this [9,28–30]. However, this means that a thermal camera needs to be placed in front of the person during at least these measurements interfering with the activity and predicting the stress may indeed occur in that period of time.

In addition, most studies on the use of physiological measurements for mental stress quantification or automated inference use a relative long term window of data (several minutes to a few hours). To achieve reliable results with a smartphone, a user has to continuously attend to the sensors (keeping the finger stable in front of the sensors and the nose within the smartphone display). This is extremely inconvenient, and greatly limits the use of this technology. There are two reasons for the requirement of a user’s continuous attention. First, imaging capabilities are affected by motion artefacts (and viewing angles). Second, environmental noises such as lighting conditions and environmental temperature [31,32] can corrupt measurements. The easiest way to address this is to require the person to remain stationary during the recording. Although algorithms could be further improved to address such issue, this would lead to increase computational cost. Finally, keeping the finger on the PPG sensors for more than 30-40 second would not be possible due to the heat from the measurement (flash light used for PPG).

To make it possible to use smartphone imaging as a mental stress measure, this paper aims to build a fast stress recognition system that only requires a short period (20 seconds) of time for the cardiovascular measurements. In particular, we aim to contribute on three fronts. First, we propose a new preprocessing step to enhance the quality of the signals (especially the thermal signals). Second, we developed a new method to address inter-subjective variability inherent to self-report of mental stress scores. Finally, we show how currently used features to extract thermal and PPG signals are low correlated with the scores and proposed to use low-level features and let the neural network learn the high level ones. We evaluate the approach on a new dataset purposely collected for this study.

## Methods

The focus of this section is on building a system that can quickly infer a person’s perceived stress using smartphone-integrated PPG and thermography.

First, we describe a recording interface and a set of techniques to produce reliable HRV parameters from the PPG sensor and sequential nose tip thermal variations from the thermal imaging sensors. We then introduce our study protocol to induce mental stress and collect 20-second sequential data of cardiovascular events and self-reports of perceived mental stress scores. Lastly, we extract low and high level features and present the performances of our system over the two sets of features and sensor modalities. We conclude by discussing by comparing our approach to data labeling with standard approaches to discuss effect of inter-subjective variability in reporting stress scores.

### Toward Smartphone as a Reliable Multiple Cardiovascular Measure

The main cardiovascular sensing channels of this work are the rear RGB camera of a mobile phone (LG Nexus 5) and an additional low-cost thermal camera (FLIR One 2G) integrated into the phone. Figure 1 shows the smartphone based physiological measurement interface. Smartphone-imaging-based PPG generally requires a user to grab the smartphone body and place his/her finger over the back camera and flash light (Figure 1a,b) – in this work, we only focus on a contact-based imaging PPG which has been fully verified in clinical studies [6,33]. Given that a normal RGB camera is only sensitive to a narrow electromagnetic spectral range of visible light in the so-called visible spectrum [31], adequate lighting is required before it can be used as a PPG sensor. Therefore, a near flash LED emission is used.

To capture a time series of raw thermal sequences, we developed the signal capturing software using FLIR One library. In consideration of a human skin property, the emissivity for thermal imaging was permanently set to 0.98 [34]. Given low-cost thermal imaging systems do not guarantee to have a consistent frame rate (i.e., temporal resolution) [31], the recording interface as shown in Figure 1c has a function to store a time stamp in each thermal image frame.

**Figure 1.**
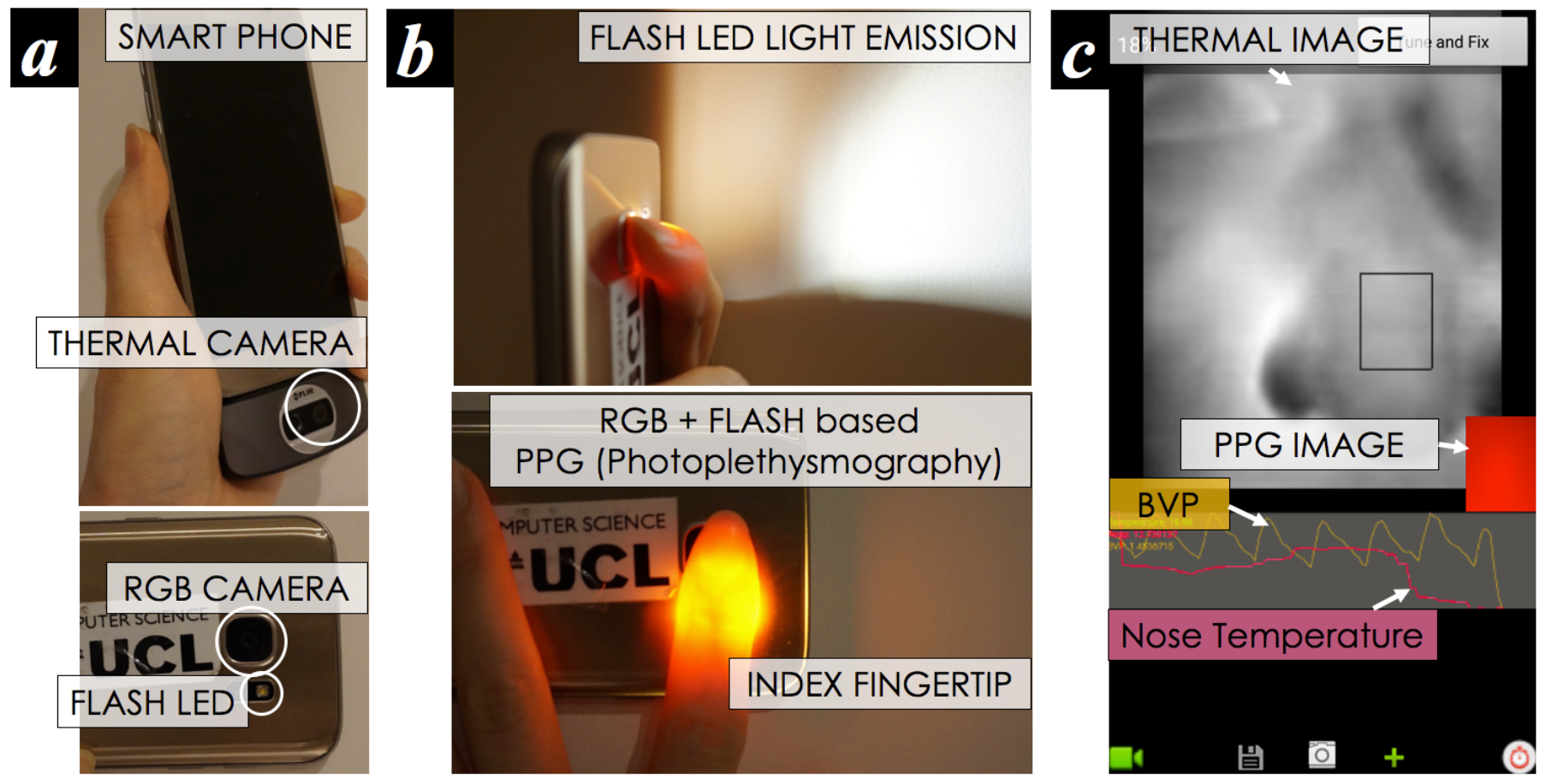
Smartphone RGB and thermal camera based physiological measurement: (a) a smartphone with an integrated thermal camera, (b) a support of flash LED emission for PPG measurement, and (c) designed interface software to collect BVP and 1D thermal signature from a facial region of interest around the nose.

#### BVP and R-R Interval Estimation through PPG Imaging

Figure 2 summarizes the approach we use to extract BVP and R-R intervals through the smartphone imaging PPG. Following [6,7,33], our method captures subtle color variations, in association with light absorptivity patterns of hemoglobin in capillaries in a person’s skin, to estimate the BVP signals. Extending the existing work, we propose and test the use of entropy patterns rather than average values of all pixels from the R channel detector (among RGB), given that averaging tends to ignore fairly small, but important variations over the color distribution [9].

**Figure 2.**
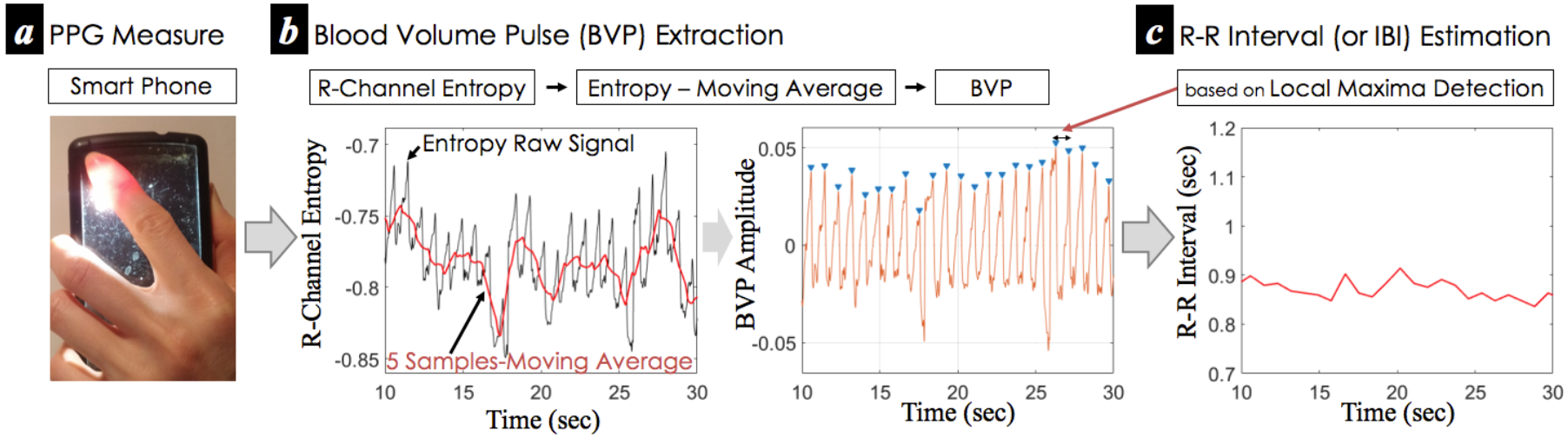
Overall procedure of BVP and R-R interval estimation from a person’s finger through the smartphone-imaging PPG.

To estimate heart rate (heartbeat), the frequency analysis method generally performs well without preprocessing the raw BVP [6,7,35]. However, given that our interest is in measuring HRV parameters from raw R-R intervals (also called Normal-to-Normal or inter-beat intervals), an additional technique is required. We introduce a technique to evenly spread each peak of BVP so as to easily use the peak (or local maxima) detection proposed in [36] to recover R-R intervals. The technique is based on the subtraction of moving average signals from the raw entropy signals (Figure 2b). For the implementation, we up-sampled the raw sequences to 256 Hz and used 5-samples moving average. Finally, we implemented a local maxima detection algorithm with a 0.5 seconds sliding window.

#### Continuous Nose Tip Temperature Measurement

As shown in the flowchart of Figure 3, the extraction of the 1D sequential nose tip thermal signature is implemented in three computational steps: i) nose-tip ROI tracking, ii) breathing dynamics removal, and iii) post processing step for extracting low thermal directionality features. In relation to the tracking steps, we take advantage of the recent advancements in thermal ROI-tracking techniques that help minimize the effects due to motion artefacts and thermal environmental changes. In particular, we used the Optimal Quantization and Thermal Gradient Flow methods (Figure 3a) introduced in [9]. Through the use of these techniques, we extract sequential average temperature values over the ROI.

**Figure 3.**
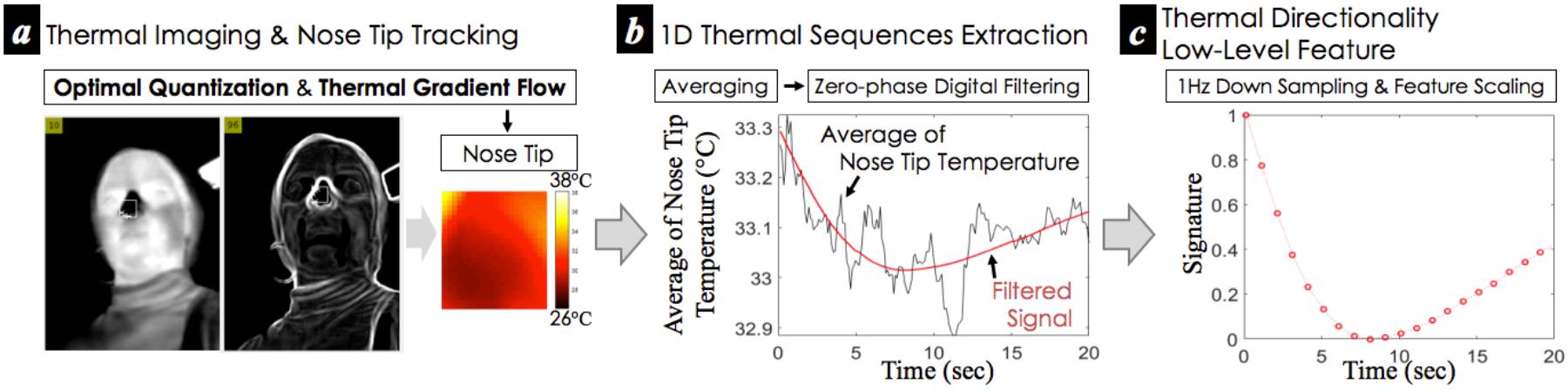
Overall procedure of the extraction of one-dimensional thermal directionality signature from a person’s nose tip through smartphone thermal imaging.

However, a remaining computational challenge is the effect due to thermal changes of the area close to the tip of the nose due to exhalation and inhalation respiratory dynamics potentially affecting the nasal temperature (see Figure 3b). To minimize the effect of these temperature changes, we propose the use of zero-phase digital filtering. For the implementation, we used a seventh order low pass digital Butterworth filter with cutoff frequency of 0.2 π (radians/sample), producing one-dimensional thermal sequences from the nasal tip. Lastly, we extract the thermal directionality signature of the nose tip which is interpolated and down-sampled to 1Hz (to get 1 second-interval low level features) to address the unsteady frame rate of the camera we use (Figure 3c). This automatic and continuous measurement of the thermal directionality signatures can address limitations of the manual analysis used in [23,26,27], in turn possibly contributing to the automatic process to infer a person’s stress.

### Data Collection Protocol

A data collection study was carried out to gather physiological data from participants during different tasks inducing different levels of mental load. The data collection protocol is defined below.

#### Participants

A total of 17 healthy adults (mean age 29.82 years, SD=12.02; 9 female) of varying ethnicities and different skin tones (pale white to black) were recruited from the University College London and non-research community. Each participant was given the information sheet and the informed consent and demographics forms prior to data acquisition. The study was conducted in a quiet lab room with no distractions and controlled ambient temperature (around 20°C). Participants were informed that they could stop the study at any time if they feel uncomfortable. Only one experimenter was present in the room during the data collection but kept not too close to the participant (longer than 1.5 m distance). The experimental protocol was approved by the Ethics Committee of University College London Interaction Centre (ID Number: STAFF/1011/005).

#### Task Structure and 20s Measurement of Lasting Stress-induced Physiological Events

We designed a stress induction study protocol to collect physiological data and subjective self-reports in association with mental stress levels due to mental load [37]. From the literature on mental stress induction studies in psychology, neuroscience and affective computing (e.g., [2,20,22,38]), we chose two cognitive-load induction tasks - the Stroop Color Word test [39] and the Mathematical Serial Subtraction test [40]. These tests were selected as they have been shown in various studies to induce mental stress and also they have been used in other thermal imaging studies. Each task was divided into two sub-tasks with different difficult levels (i.e., easy and difficult; Se – Stroop easy, Sd – Stroop difficult, Me – Math easy, Md-Math difficult) and counterbalanced in Latin squared design as in [17].

Given our focus on the use of smartphone PPG and thermal imaging which pose constraints on the length of measurement period as discussed above (in particular the PPG sensors does not allow to keep the finger on it more than 30-40 second due to a considerable amount of heat emitted from the flash LED), we gathered our measurements during a 20 second window just after each task. The aim is to capture cardiovascular measurement related to stress-induced physiological responses and their dynamics when the stressor has ended instead of measuring the signals during each task (Figure 4). Overall, this study protocol is composed of:

a. introduction, information/consent/demographics forms provided (10-20 min)
b. **sitting, resting** (5 min)
c. **20s measurement** and **self-reporting** of perceived stress (1-2 min)
d. **Stroop Test 1** (5 min)
e. **20s measurement** and **self-reporting** of perceived stress (1-2 min)
f. break (5 min)
g. **Stroop Test 2** (5 min)
h. **20s measurement** and **self-reporting** of perceived stress (1-2 min)
i. break (3 min)
j. **sitting, resting** (5 min)
k. **20s measurement** and **self-reporting** of perceived stress (1-2 min)
l. **Math Test 1** (5 min)
m. **20s measurement** and **self-reporting** of perceived stress (1-2 min)
n. break (5 min)
o. **Math Test 2** (5 min)
p. **20s measurement** and **self-reporting** of perceived stress (1-2 min)
q. break (5 min)
r. wrap-up and participant’s feedback (5-20 min)

**Figure 4.**
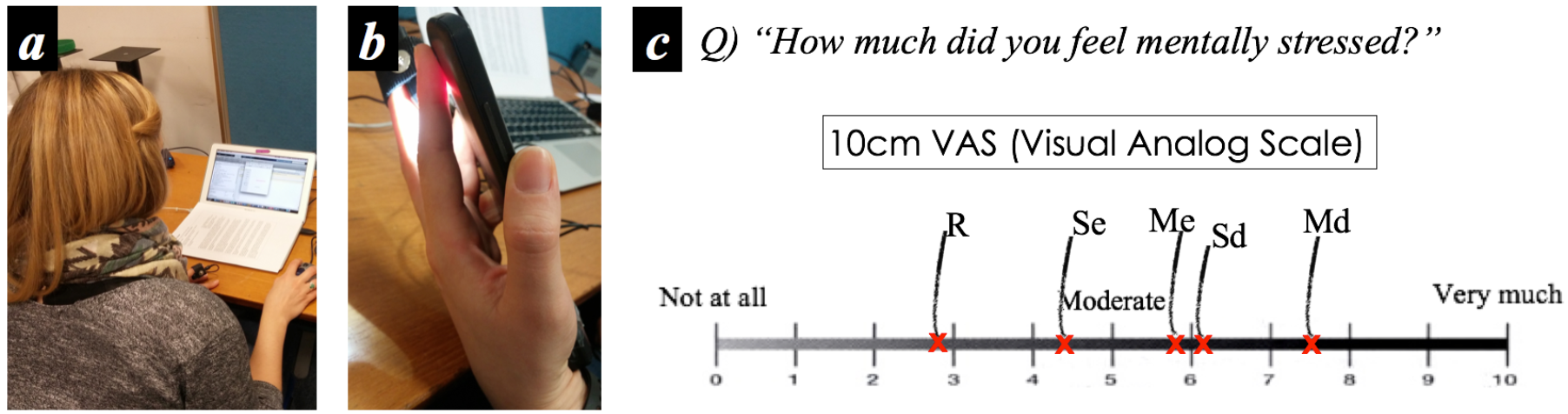
Experimental setup and self report question: (a) during each stress-induction task session, (b) 20second physiology measurement after sessions, and (c) 10cm VAS based questionnaire (R: Rest, Se: Stroop easy, Sd: Stroop difficult, Me: Math easy, Md: Math difficult).

#### Additional Stressors

However, the lack of social evaluative threat and other stressful components beyond cognitive load could be a limitation of the use of both the Stroop and math tasks for mental stress induction [2,41], although it is evident that mental stress occurs in cognitive overload [37]. Hence, we introduce further stressors: a) *social evaluative threats* (close observation and assessment of a person’s performance [2,41]), b) *time pressure* (1.5 second limitation for each Stroop question [38], and 7.5 second for each arithmetic question [41]), and c) *loud sound feedback*, in particular, an unpleasant sound for wrong answers [26].

#### Self-Report of Perceived Mental Stress

After the end of each 20 second physiological measurement, all participants were asked to answer a questionnaire about their perceived level of mental stress. We used a 10-cm Visual Analogue Scale (VAS), which allows participants to answer on an analog basis (continuous) to help avoid non-parametric properties [42]. The question asked was *“How much did you feel mentally stressed?”* (ranging from 0, not at all, to 10, very much). Only one VAS straight line was used for collecting self-reports of each perceived stress level along with each session to make participants easily remember and compare their previous scores as shown in Figure 4c. The comparison lets us combine a numerical type of approach to self-reporting with a ranking one as comparison is generally more reliable that simple quantization of a subjective state [43–45]. The labels in Figure 4c have been added to figure by the researcher to clarify their reference to each of the tasks (R: Rest, Se: Stroop easy, Sd: Stroop difficult, Me: Math easy, Md: Math difficult).

### Automatic Inference of Perceived Mental Stress from 20s Measure

#### Low-Level and High-Level Features from Cardiovascular Events

The 20 second-cardiovascular measurement with the developed interface (Figure 1c, 4b) produces the following *raw signals*:

a. one-dimensional R-R interval sequences
b. one-dimensional thermal directionality sequences

We take an array of the R-R intervals (Figure 2c) and the 1Hz-sampled thermal directionality samples (Figure 3c) as *low-level* features representing each modality throughout this paper.

To extract *high-level* HRV features from R-R intervals, we followed earlier studies on stress inference using HRV [19–22,46]. After the pre-processing method proposed above, we extracted the following hand-engineered HRV features:

a. HRV F1: LF Power
b. HRV F2: HF Power
c. HRV F3: LF/HF ratio
d. HRV F4: SDNN (standard deviation of R-R intervals)
e. HRV F5: RMSSD (root mean square of the successive differences of R-R intervals)
f. HRV F6: pNN50 (ratio of the number of the successive differences of R-R intervals greater than 50ms *of the total number of R-R intervals*) As for *high-level* features representing the nasal tip thermal signature, we propose the use of basic statistical features:
g. Thermal F1: SSDTD (sum of the successive differences of the 1Hz-sampled thermal directionality)
h. Thermal F2: SD2TD (standard deviation of the successive differences of the 1Hz-sampled thermal directionality)
i. Thermal F3: SDTD (standard deviation of the 1Hz-sampled thermal directionality)

Here, the sliding of a signal window was not applied to the feature extraction given the short period of time measures.

#### Labeling Strategy and Machine Learning Classifiers

Given the focus on the automated inference of a person’s perceived stress level, the labeling of self-reported stress scores is an important step. However, inter personal variability has been repeatedly reported on self-reports of perceived mental stress [16,17,47]. This is a key issue which must be overcome if we are to create automatic stress recognition systems that can generalize across people. Adapted from our earlier work [17], the normalized K-means clustering technique is used to label the measured events. In detail, all perceived stress scores collected from each participant are normalized through feature scaling with one’s minimum and maximum scores. Then, the k-means algorithm (k=3) is used to group three levels of perceived stress scores across participants, best corresponding to “None or low stress”, “Moderate” and “Very high” on the VAS we used (see Figure 4c). In this paper, we focus on discrimination between two levels of stress that we call for conciseness *No-stress* and *Stress* groups in accordance with the two clusters of participants’ perceived stress scores. The *No-Stress* group refers to the “None or low stress scores”. The *Stress* group refers to the combination of Moderate and very high scores obtained following the technique presented in [17] due to limited data for building a more refined model.

This labeling strategy process (denoted as L1) is used to label the measured signatures from each 20second slot.

Furthermore, we explore the possible effect of different data labeling strategies: a) L2: combining the first and second groups from the k-means (k=3) into No-stress rather than the binarization used for L1, b) L3: k-means with k=2, and lastly, c) L4: the original stress scores divided by the point between “not at all” and “moderate” (approximately, 3.334 in Figure 4c). The aim of L2 and L3 was to understand the sensitivity of our approach in separating intermediate level of stress with the other two classes. L4 was used a way to compare with the standard technique used in the field.

Two machine learning algorithms were tested. The first algorithm we use is a k-nearest neighbor classifier (denoted as kNN, k=1), using the extracted high level features listed above as in [46]. The second algorithm is a feed-forward and back-propagation based single layer neural network which uses the original input sources (i.e., 1D R-R intervals and 1Hz sampled thermal sequences) as low-level features. By choosing this second algorithm, we aim to address the limitations of the use of handcrafted features which may simplify a person’s dynamic physiological events, and in turn possibly missing some fast informative moments. In particular, in the case of instant measurement (short period of time), this cannot be compensated by the use of a sliding window producing sequential feature values (e.g., a 120 seconds sliding window used in [22] to continuously produce HRV features during a 180 second task session). The use of artificial neural networks can empower automatic learning of informative physiological features [17] by using the back-propagation algorithm to repeatedly tune internal parameters of a machine learning model to better learn features (this is also called representation learning).

For the implementation of the neural network, we tested two different hidden layer node sizes: a) 80 (smaller, denoted as NN1) and b) 260 (larger, NN2). A mean and standard deviation of z-scores of training data were used to normalize both the training and testing data. The sigmoid was used as activation function. In the training process, a fixed learning rate of 0.5 was used for 100 epochs. Lastly, feature scaling was applied to the case of the low-level thermal directionality features so as to minimize different overall temperature levels across participants and sessions (Figure 3c). From our preliminary observations, the temperature difference between the lab room and the place each participant stayed before existed. This may lead to wrong data collection by capturing a thermal direction due to the room temperature at the start of the experiment. Hence, we discarded the data collected in the beginning. Hence, we mainly use the physiological data from the second resting period to represent the resting condition.

## Results

In this section we evaluate our proposed approach. First, we report the analysis of the collected data. Second, we discuss the recognition performance of our system over the different modalities and levels of features. Finally, we compare the results over the different labeling approaches.

### Signal Quality of Measured Physiological Patterns

First of all, we tested the physiological measurement reliability. From the 17 participants, we collected 82 sets of R-R intervals from 20 seconds instant measurements after the end of both the two Stroop and two Math tasks and after the resting session. This gives rise to 85 possible sets. However, 2 sets of recordings had to be discarded due to phone battery issues at the end of the study, and 1 set was not recorded as a participant clicked the turn-off button on the phone by mistake. As for the thermal data, in addition to the 2 sets listed above, 6 further sets had to be rejected because of participants’ big movements (coughing severely) and hiding their noses. The remaining data consisted of 77 recordings.

Given the cardiac measurement ability of smartphone PPG thoroughly verified in earlier studies (e.g., [6,33]), we only tested the reliability of cardiac pulse signals measured by our approach by using a periodic signal goodness metric based signal quality index (SQI) [9,35]. The SQI is used by setting a frequency range of interest of physiological signals we are interested in [35]. Figure 5 shows the cardiac pulse rate SQI distributions of the measured signals. It shows that the measurement was generally reliable (mean of *Pr*=0.9584, SD=0.0151).

**Figure 5.**
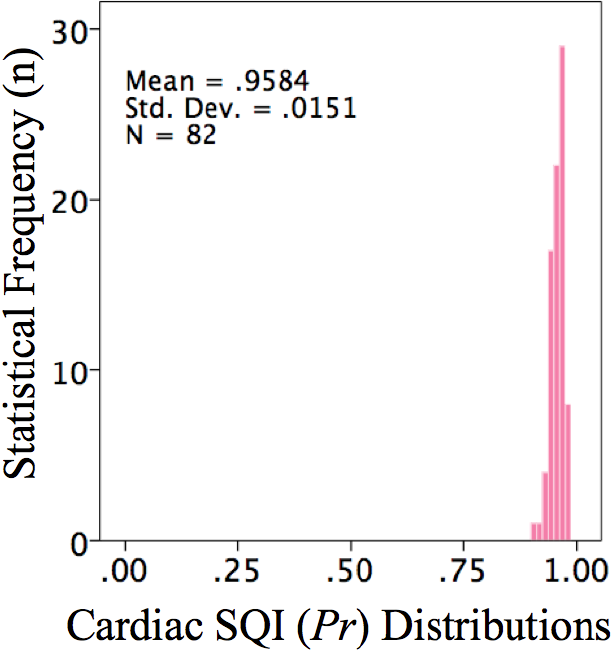
Cardiac pulse rate SQI distributions of the collected data through smartphone PPG.

Figure 6a shows examples of thermal images taken from the participants during the data collection study. From our preliminary and empirical observations, we found that respiration may influence the nasal tip temperature as discussed in the previous section. For example, along with inhalation volume, the nose tip surface becomes colder as shown in Figure 6b. This was due to the colder air (i.e., the air in the room was colder than a person’s body) flowing into the nostril during the inhalation. Hence, we tested the effect of respiratory activity on the nose tip temperature by using the goodness probability with the respiratory rate of interest and found the participant’s exhalation and inhalation cycled events affected the nose tip temperature measurements (M=0.8998, SD=0.0620), producing less reliable (i.e., representing stress) thermal directionality signatures. This suggests that a filtering technique is needed to avoid wrong classification (see Figure3b). Previous studies have not considered this issue possibly because they only relied on using start and end measurements and did not use information about the fluctuation of the profile.

**Figure 6.**
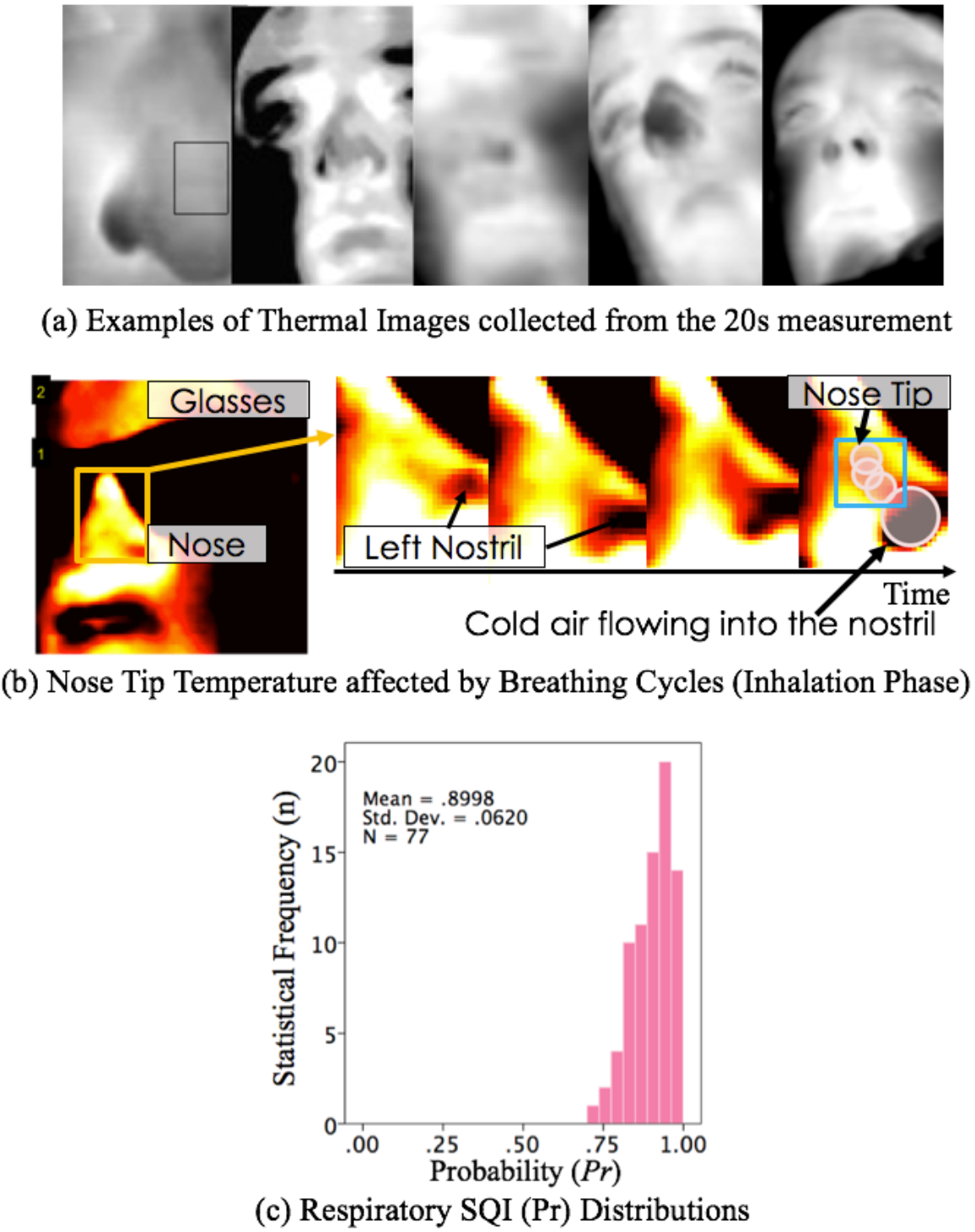
A person’s respiratory cycles may deteriorate the reliability of inferring the nasal tip temperature: (a) examples of thermal images from participants (view angles were differently set during the measurement), (b) empirical observation (the nasal tip surface became colder (yellow: warmer, red: moderate, black: colder) along with during inhalation - and warmer but not shown here during exhalation), and (c) respiratory SQI distributions of the collected data through smartphone thermal imaging.

### Self-Reported Stress Ratings and High-level features

Another important step was normalization of the self-report of stress scores. Given the limited set of data to carry a multi-level model and the subjectivity of stress ratings, in our study we focused on binary classification of stress: no/low stress vs medium/high (or very high) stress. Figure 7 shows the overall distributions of selfreported perceived stress scores from the 17 participants over the sessions, including the resting period. The raw self-reported scores seemed to be evenly distributed as can be seen in Figure 7a. However, the distribution is not ideal to cluster their values into two groups (some drops were due to a quantization effect in the data visualization). Interestingly, the within participant normalization technique discussed in the data collection protocol session instead transformed the distribution of the score into a bimodal distribution where two big peaks (mode: most frequently occurring value) exist (Figure 7b). The K-means based labelling was able to cluster the normalized perceived scores into two groups (see the line in Figure 7b).

**Figure 7.**
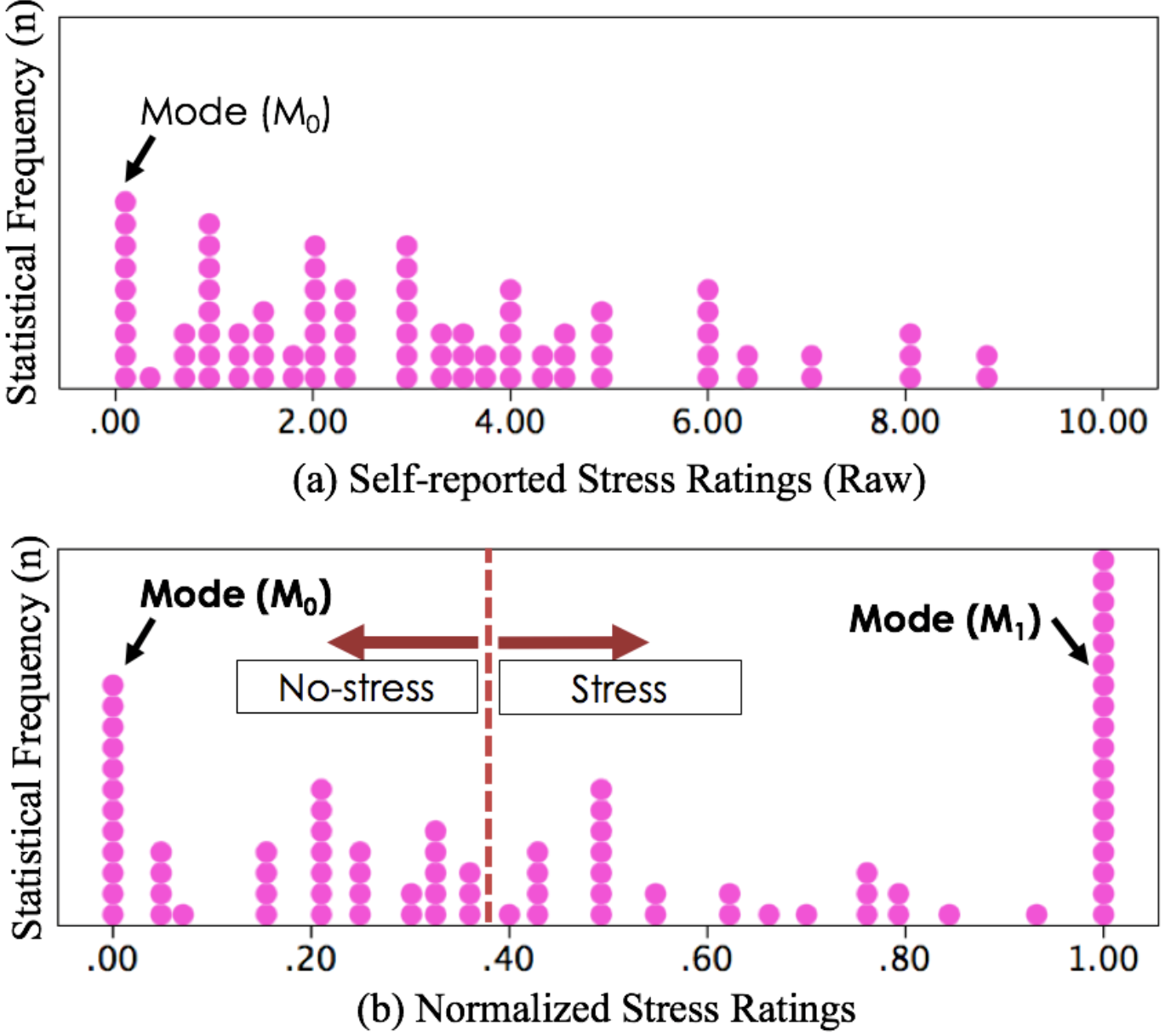
Overall self-reported stress score distributions (from 17 participants over the sessions including the resting period): (a) original scores, (b) normalized stress scores (normalization of scores from each participant) and K-means based grouping (No-stress and Stress groups)

**Table 1.**
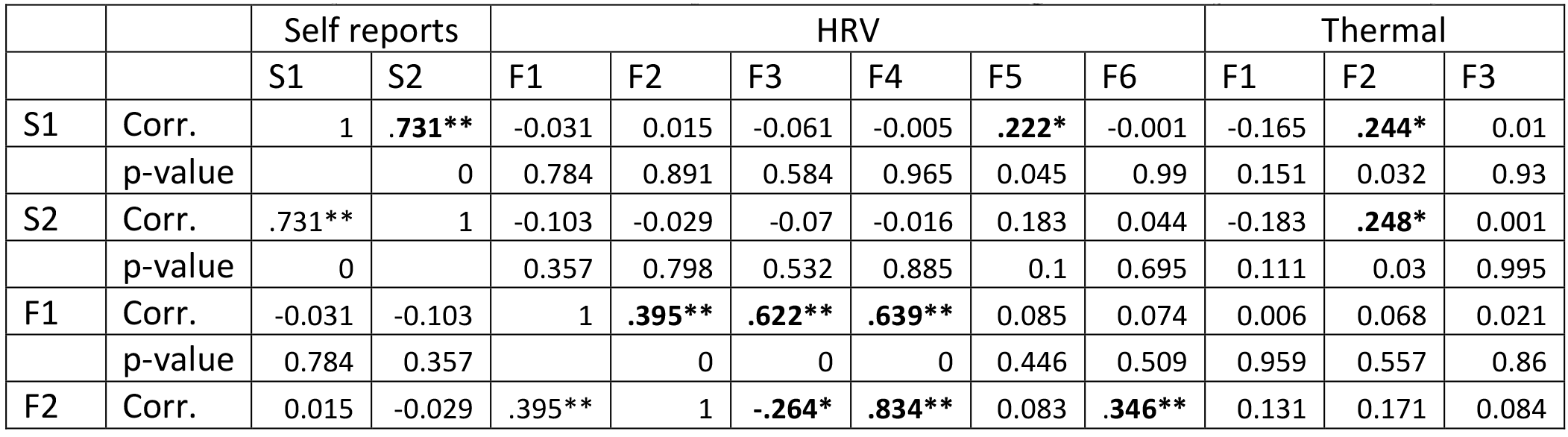
Pearson correlation coefficients across self reports, HRV and thermal high-level features (S1=Normalized self reported scores, S2=original self reported scores)

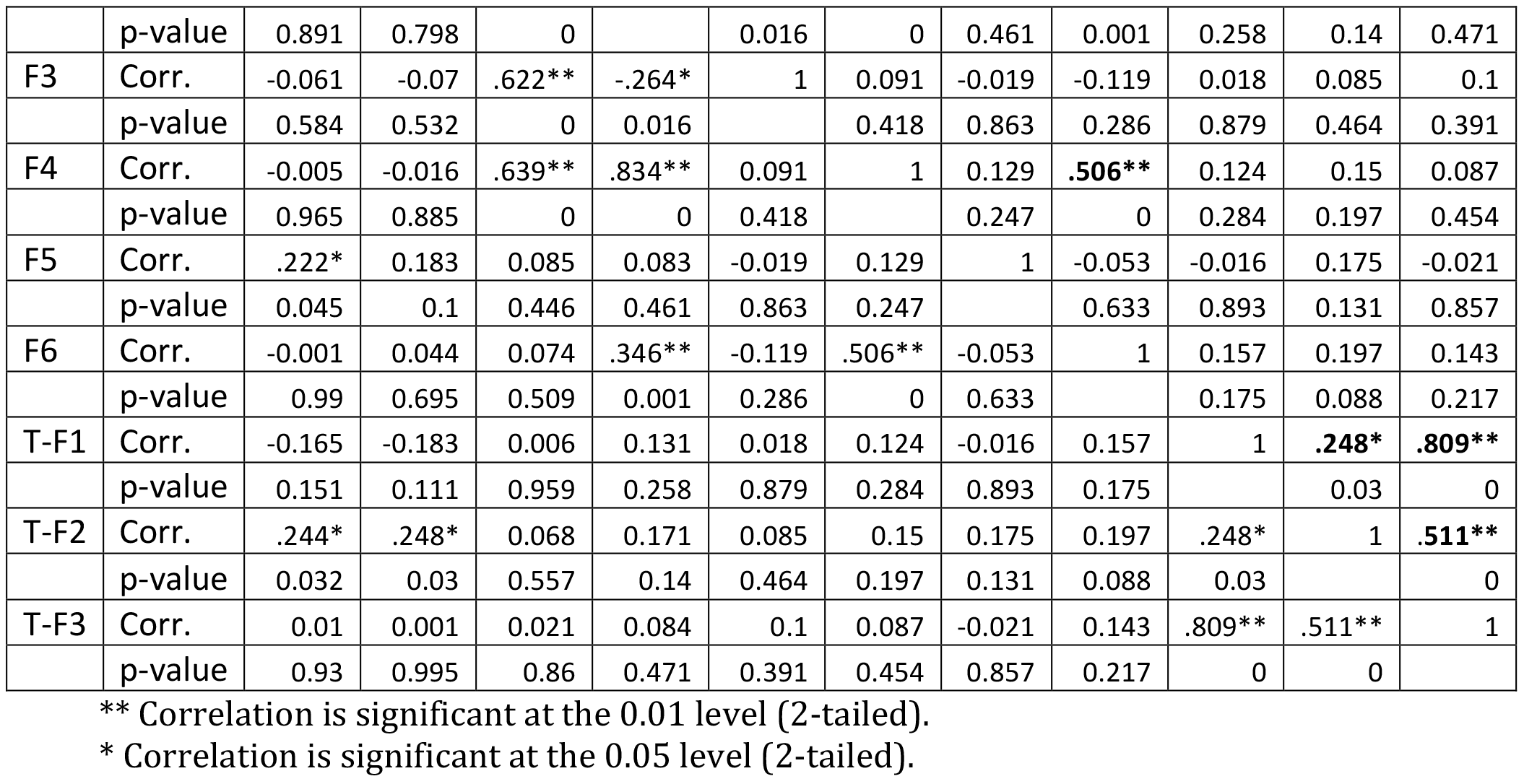

We tested the correlations between the original self-reported scores, normalized self-reported scores and high-level features extracted from the HRV and thermal signatures as summarized in Table 1. The normalized self-scores maintained a high correlation with the original scores (r=0.731, p<0.001). While some features from each physiological sensing channel were significantly correlated between themselves (e.g., HRV F2 and F4: r=0.834, p<0.001; Thermal F1 and F3: r= 0.809, p<0.001), the correlation values were lower across sensing channels. In addition, each hand-engineered high-level feature (6 from HRV and 3 from thermal directionality) was less correlated with self-reported stress ratings (e.g., Thermal F2 – original scores: r≤0.248, p=0.03), indicating that each feature alone was not sufficient to lead to high discrimination between levels of stress.

Figure 8 shows the general trends of each feature value distribution along with sessions (rest and four stressful events, i.e., Stroop easy/difficult and Math easy/difficult) and labels produced by the labelling technique. Although the features responded to each session event differently (Figure 8a), their patterns were irregular over sessions. For example, Thermal F1 appeared to strongly decrease during the *Math difficult task*, Thermal F2 increased with the *Stroop difficult task*, but less during the *Math difficult task.* While HRV F5 was generally high after both Math easy and difficult task sessions, Thermal F2 was generally high after when the Stroop difficult and Math difficult tasks were completed. This lack of a clear pattern may indicate that each feature itself is unclear about the stressful event and the cognitive load and mental stress may be tangled together, leading to this ambiguity. This may be a reflection of what different processes are at work. In the case of the features patterns along with the perceived stress label, Thermal F1 was generally lower from the stress group than that from no-stress group – this is consistent with findings from literature (i.e., decrease in the nasal tip temperature in association with mental stressors [23,26,27]).

**Figure 8.**
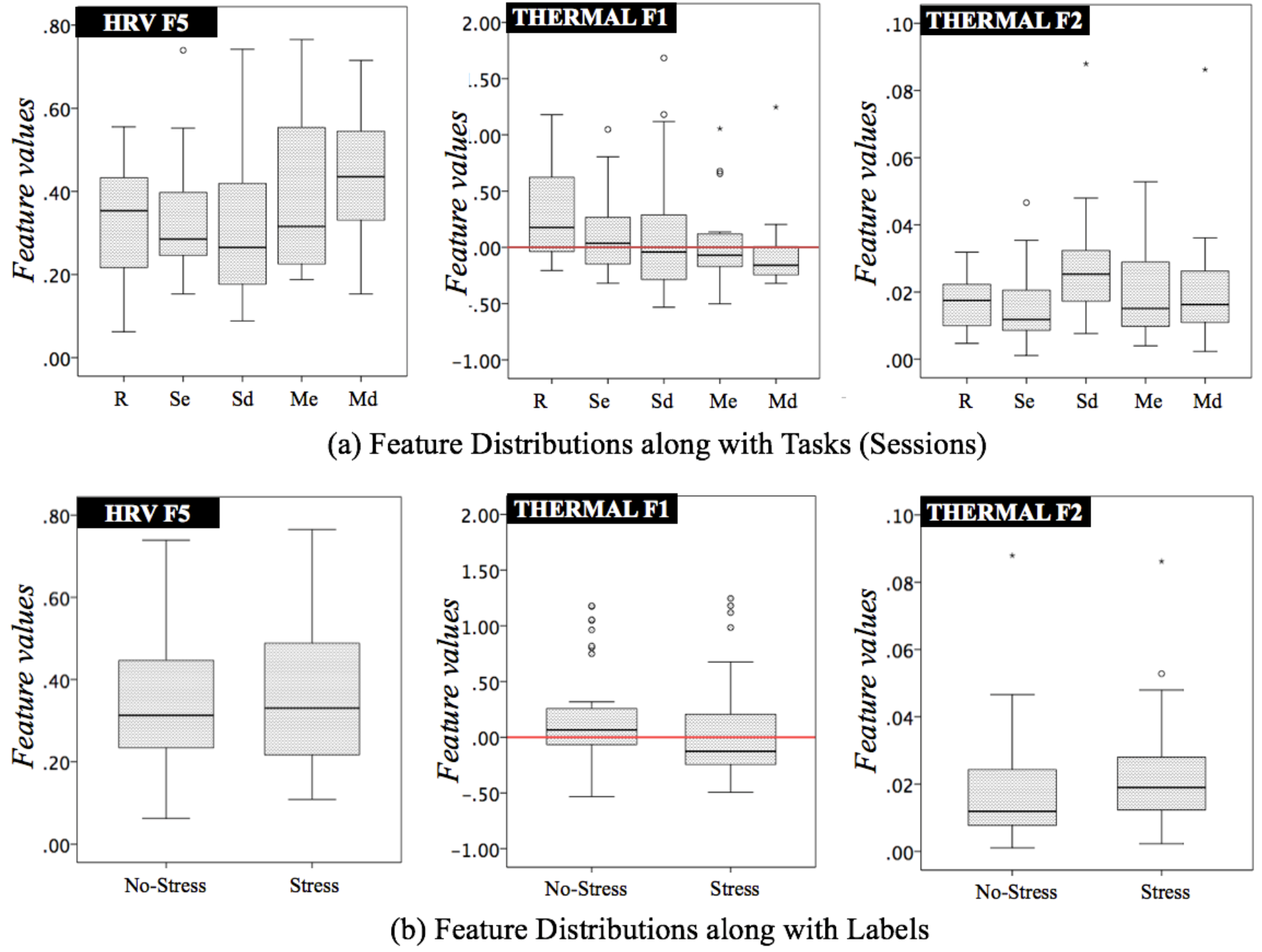
Bar plots of 95% confidence intervals in each feature value distribution over (a) each session (R: Rest, Se: Stroop easy, Sd: Stroop difficult, Me: Math easy, Md: Math difficult) and (b) label produced by our labelling technique. The three features (HRV F5 - root mean square of the successive differences of R-R intervals, Thermal F1 - sum of the successive differences of the 1Hz-sampled thermal directionality, F2 - standard deviation of the successive differences of the 1Hz-sampled thermal directionality) which were best correlated with self-reported scores are used.

### Instant Stress Inference Results

To evaluate the performance of instant stress recognition, we used a 17-fold leave-one-subject (participant)-out (LOSO) cross-validation. We also used this to test the ability to generalize to unseen participants (*one size fits all*) [17,47]. Figure 9 summarizes the accuracy results of the three classifiers (NN1, NN2, kNN) from the LOSO (N=17) for three different cases: a) multimodal-approach by simply combining features from both sensing channels (HRV, Thermal), b) unimodal approach using thermal features, and c) unimodal approach using HRV features. Both neural networks NN1 and NN2 only used low-level features (i.e., an array of the R-R intervals for HRV and the 1Hz-sampled thermal directionality samples for Thermal).

Overall, the NN2-based multimodal approach produced the highest accuracy of 76.96% in discriminating between no-stress and perceived stress. The false positive and false negative cases were also much lower than other cases (see Figure 9b). The NN1, whose hidden layer is smaller than that for NN2, produced the second highest accuracy (76.47%). The similar pattern was shown from the case of thermal-based unimodal stress inference. Interestingly, the kNN which is based on the use of high-level features (i.e., hand crafted 6 HRV and 3 thermal features) performed better for HRV unimodal inference than the other approaches. This might be due to the lack of feature weighting or feature selection phases in this work. Nonetheless, the ability of the artificial neural networks in the automatic feature learning and inference of perceived stress levels is remarkable as successful NN-based deep learning approaches in other applications [48].

**Figure 9.**
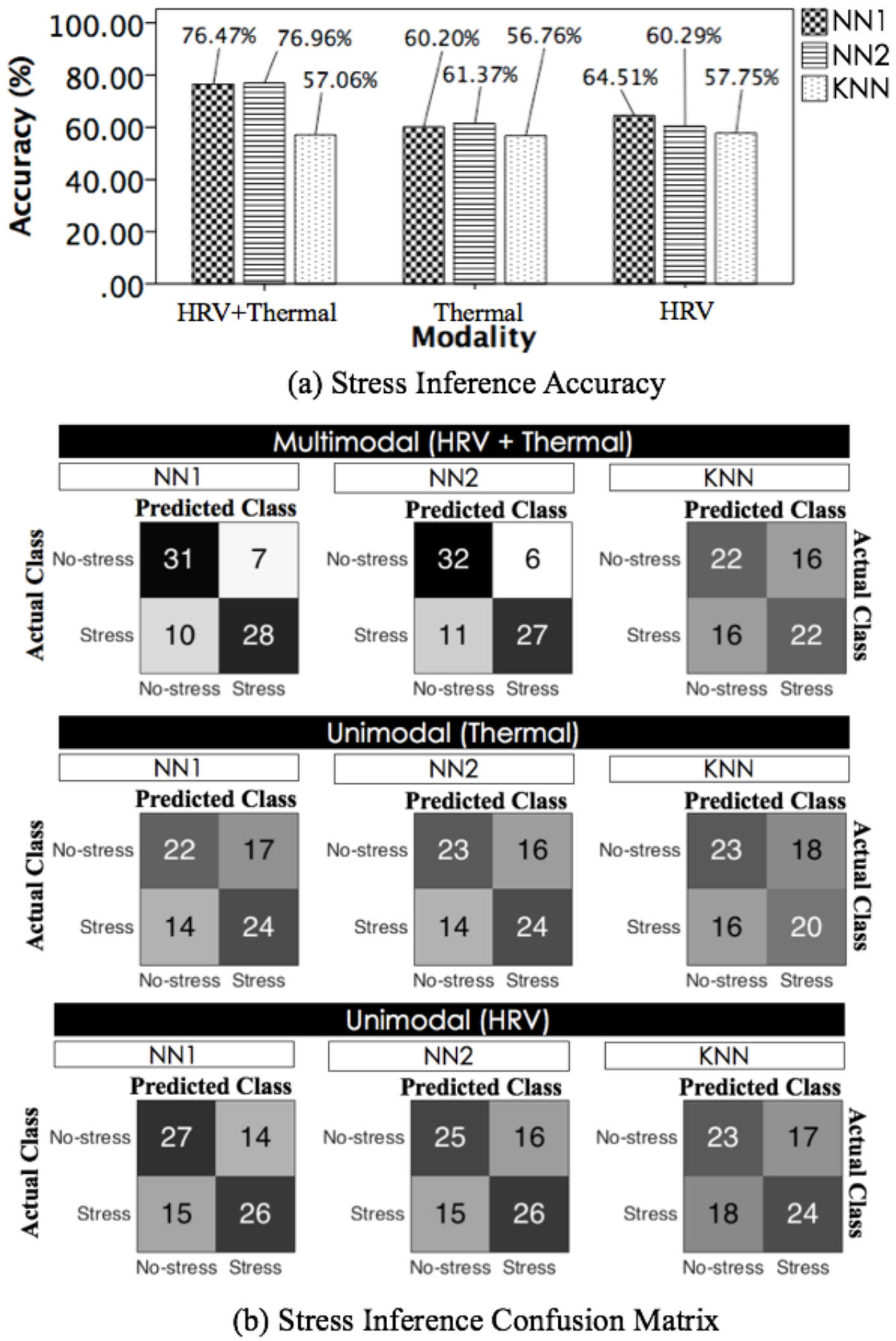
Summary of (a) inference accuracy and (b) confusion matrix achieved from three classifiers NN1, NN2 and kNN along with modality (Multimodal: HRV+Thermal, Unimodal: Thermal, HRV) (each number means the number of cases).

Lastly, we investigate the effect of the self-reported score normalization and k-means clustering in better labeling the perceived stress levels. We evaluated the pe formance of the built inference systems (with the multimodal features) when using different labelling strategies (L2-L4, introduced in the previous subsection ‘labeling strategy’). Figure 10 summarizes the accuracy results for four different strategies - L1: proposed method (Figure 7b), L2: k-means with k=3, but combining no-stress and moderate level stress scores as a same group, L3: k-means with k=2, dissecting the moderate level scores into no-stress and stress, and L4: original scores divided by a point between no-stress and moderate levels (i.e., 3.334 of 10, see Figure 4c). Overall, all three classifiers obtained the best accuracy for L1 and the worst performances for L3 and L4.

**Figure 10.**
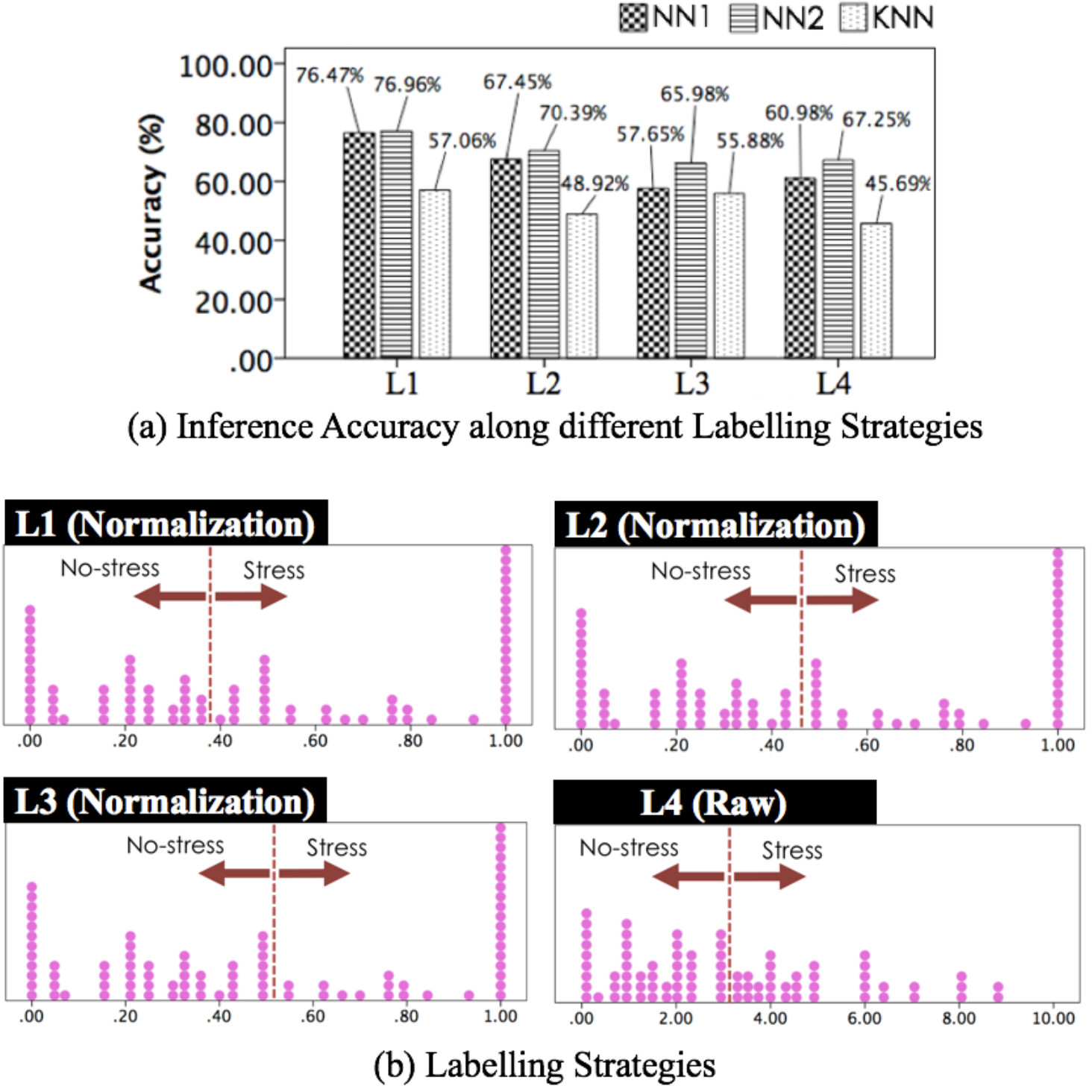
Summary of (a) inference accuracy along with (b) different labelling approaches (L1: K-means with K=3 and combining moderate and high stress scores, L2: K-means with K=3 and combining no-stress and moderate level stress scores, L3: K-means with K=2, L4: original scores divided by the border between no-stress and moderate levels).

Rather than adjusting the labeling to obtained the best performance, our aim was to understand how the normalization of the labelling to address class separation and inter-person variability in subjective self-reports may lead to more effective modeling. Indeed, in the literature approaches such as L4 or to a certain extent L3 are generally used limiting the ability to generalize across participants ([43,49]). We were also interested to understand how sensitive the system was in separating more levels of stress by using the same dataset and merging the intermediate levels with one of the two classes (L1 and L2). The results show that more data are needed to separate intermediate level of stress from high level of stress. suggesting that the labelling method L1 is capable of better separating the bimodal distribution of normalized self-reported scores and seems to better address the inter-personal variability issue.

## Discussion

The overall results of the quantitative analysis and classification tasks show the possibility of using a smartphone-based biomedical imaging capabilities to build an automated stress recognition system that could be used as an input to stress management support apps.

The first contribution made is in the way the signals are pre-processed to enable robust measurements over short periods of time. Through the data collection from the designed stress induction study, we first evaluated the ability of the built cardiovascular monitoring algorithms in enabling a smartphone to reliably monitor a person’s HRV and thermal propagation events. In particular, we found that a person’s respiratory cycles interfere with accurately capturing the thermal directional variation of a person’s nasal area, an area informative of one’s mental stress (Figure 6). Hence, a new thermal directionality temperature signature was designed to minimize such effect. This was achieved by the use of an advanced thermal ROI tracking and filtering techniques to remove breathing contributions (Figure 3 and [9]) of the thermal image of the nose area. First of all, this tracking made it possible to automatically and continuously monitor a subtle thermal directional variation during the measurement period. This approach is more informative than the typical approach of using a single discrete number representing the difference between two end-points. Indeed, rather than showing a clear increase or decrease, the nasal tip temperature fluctuates during such stressful tasks and its fluctuation pattern is an important source of information (e.g., suddenly decreasing and smoothly increasing in Figure 3c). However, other important issues such as person movement, covering of the nose due to coughing, perspective of the head position, etc. can contribute to reduce the quality of the signal [31,32] (e.g., P14’s coughing observed during the study), disappearance beyond a field of view (e.g., coughing and turning one’s head), and hiding of the relevant ROI (e.g., P5, P11, P14’s adjusting of their glasses hiding it behind their hand). Hence, in addition to tracking and filtering the signal, our approach aimed to keep the measuring time needed to compute stress measurement very short compared to the minimum of 2-5 minutes period considered necessary in the literature, for addressing the limitation of the long term use of smartphone biomedical imaging capabilities.

The second contribution, is in investigating the contribution of each of the signals and the features to the modeling process. The overall accuracy results in discriminating perceived mental stress levels demonstrate a smartphone-based biomedical imaging can be an effective instant stress recognizer (best accuracy: 76.96% from the 17-fold LOSO cross-validation, see Figure 9). Compared to perceived stress recognition performances of existing studies achieved from a long term measurement (up to around 70 - 80% from LOSO [47]), our method reached state-of-the-art by relying on very short period of measurements at the end of each session (data from resting period and four stressful tasks were used).

The contribution of the hand-crafted features used in the literature to the stress quantification from the instant measurement was weak (see Table 1). Given their limited correlation with stress and low performances obtained by the classifier built on them, we turned to automatic feature learning and classification using artificial neural networks (NN1, NN2). Inputting the combined low-level features from HRV and thermal signatures to a neural network makes it possible to reach higher and state of-the-art comparable good performances over the short period of time after the stressing event had finished.

Finally, we also explored how different labeling technique may affect the modeling process. How to use self-reports to label the data is a critical issue in the field due to their subjectivity. The inter-personal variability has been repeatedly reported as a main barrier for a stress inference or quantification system to be able generalize across people [16,47]. We proposed to address this problem through a two-step process. First, we transform the perceived scores so that their distribution shows bimodal properties (see Figure 7b) despite maintaining high correlation with the original scores. Second, we use a machine learning clustering technique such as k-means to better separate the scores into two classes. The comparison of the performances of the system trained with our labelling technique against most standard ones (e.g., a mid-or low-score value dividing original scores into two groups [49]) brings validity to the process (see Figure 10).

## Limitations and Future Directions

Despite the findings summarized above, there is still space for improvement toward building a reliable mental stress system that can be used by everybody in everyday situations. First, the built system only worked on binary discrimination. We tested the system on three classes of stress, but the results were at chance level. This might be due to the limited size of the dataset, especially for the medium level of stress. Deploying the built software could be a solution to build a larger dataset. With a function to collect a person’s perceived stress score (e.g., digitalized VAS sliding bar in an app), the data collection in the wild would produce a sufficient size of cardiovascular signals sets to support multi levels inference capabilities. In addition, it would be interesting to investigate how the transformation of the scores can be used to support multi-class classification.

A second limitation is a lack of a distinct baseline in our approach. We avoided the use of a baseline in the approach as in everyday life, such baseline may be difficult to detect. Even within our data, despite the resting period, participants’ scores did not show a clear no stress situation. This could be due to a lab effect but also to the multiple number of tasks used with different stress levels and the difficulty to selfreport differences of mental stress in such situations (extended deskwork in offices or during studies). Hence, rather than the use of a baseline, data collection in real-life situation and more interesting self-report methods could be used to provide more reliable scores that are able to capture subtler differences.

Lastly, this work did not include a certain level of physical activities beyond the used sedentary desk activity. It has been shown that physical activities influence a person’s cardiovascular events and the stress inference performance (e.g., [20]). Given this problem, it would be interesting to test the instant stress inference ability of our system in situations where there is a considerable amount of physical activities. At the moment, we are indeed extending our study to monitor stress in the context of industry factory workfloor.

## Conclusions

Toward building an automatic stress inference ability in HCI, this paper focuses on the use of smartphone-based biomedical imaging capabilities: imaging PPG and thermal imaging. To overcome difficulties of using smartphone imaging for long period measurements (e.g., requiring a user’s high level attention to avoid motion and illumination artefacts), we propose a novel method that requires 20 seconds to infer a person’s perceived stress level from instant physiological measurement. This is achieved by i) developing a more reliable sensing techniques to extract multiple physiological patterns (i.e., HRV and thermal directionality); ii) building an automatic feature learning-based multimodal stress recognizer; and iii) by transforming self-report scores to take into account the subjectivity of the selfreport and ensure clear separation between the level of stress to be modelled. . Through a stress induction study with 17 participants and a series of different level of stress inducing tasks, we demonstrated how this system was able to perform at state of the art level but using only 20 seconds rather than the minimum of 2 to 5 minutes of recordings. We also showed the effectiveness of multimodal fusion for producing better performances for this fast automated inference of a person’s mental stress levels. This work makes smartphone biomedical imaging capabilities more feasible for real-life applications opening new possibility for the development of mental-stress support apps and research.

## Acknowledgements

We thank all the participants participated in our experiment. Youngjun Cho was supported by University College London Overseas Research Scholarship (UCL-ORS) awarded to top quality international postgraduate students.

## Abbreviations

PPG: photoplethysmography
BVP: blood volume pulse
VAS: visual analogue scale
NN: artificial neural network
kNN: k nearest neighbor

